# Intermittent Fasting Mitigates Vascular and Neuronal Pathologies in a Mouse Model of Vascular Dementia

**DOI:** 10.1101/2022.02.22.481534

**Authors:** Vismitha Rajeev, David Y. Fann, Quynh Nhu Dinh, Hyun Ah Kim, T. Michael De Silva, Dong-Gyu Jo, Grant R. Drummond, Christopher G. Sobey, Mitchell K.P. Lai, Christopher Li-Hsian Chen, Thiruma V. Arumugam

**Affiliations:** Memory Aging and Cognition Centre, Department of Pharmacology, Yong Loo Lin School of Medicine, National University of Singapore, Singapore; Department of Biochemistry, Yong Loo Lin School of Medicine, National University of Singapore, Singapore; Healthy Longevity Translational Research Program, Yong Loo Lin School of Medicine, National University of Singapore, Singapore; Centre for Healthy Longevity, National University Health System (NUHS), Singapore; Centre for Cardiovascular Biology and Disease Research, Department of Physiology, Anatomy and Microbiology, School of Life Sciences, La Trobe University, Bundoora, VIC, Australia; School of Pharmacy, Sungkyunkwan University, Suwon, Republic of Korea

**Keywords:** Intermittent fasting, Vascular dementia, Blood-brain barrier, Chronic cerebral hypoperfusion, White matter lesions, Neuronal death

## Abstract

Chronic cerebral hypoperfusion (CCH) is an important pathophysiological mechanism of vascular cognitive impairment (VCI). The heterogeneous effects of CCH complicate establishing single target therapies against VCI and its more severe form, vascular dementia (VaD). Intermittent fasting (IF) has multiple targets and is neuroprotective across a range of disease conditions including stroke, but its effects against CCH-induced neurovascular pathologies remain to be elucidated. We therefore assessed the effect of IF against CCH-associated neurovascular pathologies and investigated underlying mechanisms. Male C57BL/6 mice were subjected to either ad libitum feeding (AL) or IF (16 hours of fasting per day) for 4 months. In both groups, CCH was experimentally induced by the bilateral common carotid artery stenosis (BCAS) method. Sham operated groups were used as controls. Measures of leaky microvessels, blood brain barrier (BBB) permeability, protein expression of tight junctions, extracellular matrix components and white matter changes were determined to investigate the effect of IF against CCH-induced neurovascular pathologies. IF alleviated CCH-induced neurovascular pathologies by reducing the number of leaky microvessels, BBB breakdown, loss of tight junctional proteins and vascular endothelial growth factors. In addition, IF mitigated the severity of white matter lesions, maintained myelin basic protein levels, while concurrently reducing hippocampal neuronal cell death. Furthermore, IF reduced CCH-induced increase in levels of matrix metalloproteinase (MMP)-2 and its upstream activator MT1-MMP, which are involved in the breakdown of the extracellular matrix that is a core component of the BBB. Additionally, we observed that IF reduced CCH-induced increase in the oxidative stress marker malondialdehyde, and increased antioxidant markers glutathione and superoxide dismutase. Combined, our data suggests that IF attenuates neurovascular damage, metalloproteinase and oxidative stress-associated pathways, and cell death in the brain following CCH in a mouse model of VCI. Although IF has yet to be assessed in human patients with VaD, our data suggest that IF may be an effective means of preventing the onset or suppressing the development of neurovascular pathologies in VCI and VaD.

## Introduction

Vascular cognitive impairment (VCI) embodies a spectrum of cognitive deficits that range from mild cognitive impairment to vascular dementia (VaD). Due to a considerable increase in the aging population, VCI is becoming a major public health concern worldwide (Rizzi et al., 2014; Roman et al., 2004). VCI is associated with cerebrovascular diseases that arise from vascular pathological processes such as atherosclerosis, microvascular protein deposits, haemorrhages and microbleeds (Akinyemi et al., 2013; Cheng et al., 2020; Gyanwali et al., 2019; Hilal et al., 2017; Skrobot et al., 2018; Xu et al., 2018). These vascular pathologies lead to a state of reduced blood flow to the brain that is referred to as chronic cerebral hypoperfusion (CCH) (Deng et al., 2018; O’Sullivan et al., 2002; Ruitenberg et al., 2005; Safouris et al., 2015; Schuff et al., 2009). Decreased cerebral perfusion has been reported to correlate with dementia severity, and shown to be a predictive marker to identify individuals with mild cognitive impairment who develop dementia (Alsop et al., 2010; Chao et al., 2010). VaD patients have neurovascular pathologies such as blood brain barrier (BBB) dysfunction, vascular damage, white matter lesion (WML) formation, glial activation, neuronal loss, and hippocampal atrophy (Muñoz Maniega et al., 2017; Schmidt et al., 2015; Wardlaw et al., 2013). The bilateral common carotid artery stenosis (BCAS) mouse model of VCI is based on ivnducing brain CCH, and is a well established model for the neurovascular pathology observed in VaD patients (Madureira et al., 2011; Shibata et al., 2004).

CCH induces a cascade of cellular and molecular mechanisms that contributes to the pathogenesis of VCI – including oxidative stress and inflammation. CCH has been reported to increase the levels of matrix metalloproteinases (MMPs), inflammatory cytokines such as interleukin 1 beta (IL-1β), interleukin 6 (IL-6) and tumour necrosis factor (TNF) (Belkhelfa et al., 2018; Ceulemans et al., 2010; Kim et al., 2018; Nakaji et al., 2006; Rea et al., 2018; Schmitz et al., 2015; Zuliani et al., 2007) and promote cortical microbleeds, which are structural lesions in the brain that compromise cerebrovascular integrity (Okamoto et al., 2012). A high frequency of microbleeds have been associated with an increased risk of cognitive deterioration and dementia (Akoudad et al., 2016; Lee et al., 2018; Nakamori et al., 2020). While microbleeds in the brain account for gross pathology in CCH, at the cellular level, the structural and functional integrity of the brain depends on the delivery of substrates between the blood and the brain through the blood brain barrier (BBB). Indeed, it has recently been reported that BBB dysfunction may be an initiator of WML and cognitive decline in VaD (Huang et al., 2018; Wardlaw et al., 2017; Zhang et. al., 2018).

Intermittent fasting (IF) is defined as an eating pattern that cycles between periods of eating and fasting. IF has been extensively reported to extend lifespan and decrease the development of age-related disorders including cardiovascular, metabolic and neurodegenerative diseases (Mattson, 2000). Recently, IF has gained much interest as being more effective than caloric restriction for inducing neuroprotective effects in the brain (Dias et al., 2021; Pak et al., 2021). Mechanistically, IF has been reported to enhance neuroprotection through an upregulation of neuroprotective proteins while reducing the activation of pathological pathways involving cellular stress, inflammasomes, and programmed cell death under ischemic conditions (Ahn et al., 2018; Arumugam et al., 2010; Fann et al., 2014; Mattson, 2005; Poh et al., 2021; Stranahan & Mattson, 2012). However, the effects of IF on neurovascular pathology during CCH has not been studied.

In this study, we demonstrate for the first time that IF promotes neuroprotective effects in a BCAS model of VaD by maintaining the integrity of the neurovascular structures in the brain. We specifically show that IF attenuated vascular pathology by reducing microvascular leakage and BBB dysfunction, while maintaining expression of tight junction (TJ) proteins and vascular growth factors. IF was also effective in decreasing WML formation, hippocampal neuronal cell death and cell death markers, while maintaining myelin basic protein levels. Our data suggest that the effects of IF on the structural integrity of the neurovasculature may be mediated through mechanisms that decrease oxidative stress and matrix metalloproteinase expression. Overall, our findings indicate that IF may be a potential therapy in reducing and preventing vascular pathology associated with VaD.

## Materials and Methods

### Experimental Animals

Six-week-old wild-type male C57BL/6NTac mice (n=150) were obtained from InVivos, Singapore and housed upon arrival at the National University of Singapore Animal Facility. As shown in **Figure 1A**, at eight weeks of age, the mice were randomly assigned to either the *ad libitum* (AL) or the intermittent fasting (IF) diet regimen. Mice under the IF regimen underwent 16 hours of fasting every 24-hour period, with food available for the remaining 8 hours from 07:00 to 15:00 for four months (Lights on at 7:00, lights off at 18:00). All in vivo experimental procedures were approved by the National University of Singapore, Singapore Animal Care and Use Committee and performed according to the guidelines set forth by the National Advisory Committee for Laboratory Animal Research, Singapore. All efforts were made to minimize any suffering and numbers of animals used. All sections of the manuscript were performed and reported in accordance to ARRIVE (Animal Research Reporting In Vivo Experiments) guidelines. In a separate set of experiments, animals from both AL and IF groups (10-20 animals per group) underwent physiological measurements such as body weight, blood glucose and ketone levels throughout the four months as shown in **Figure 1B-D**.

**Figure 1:**
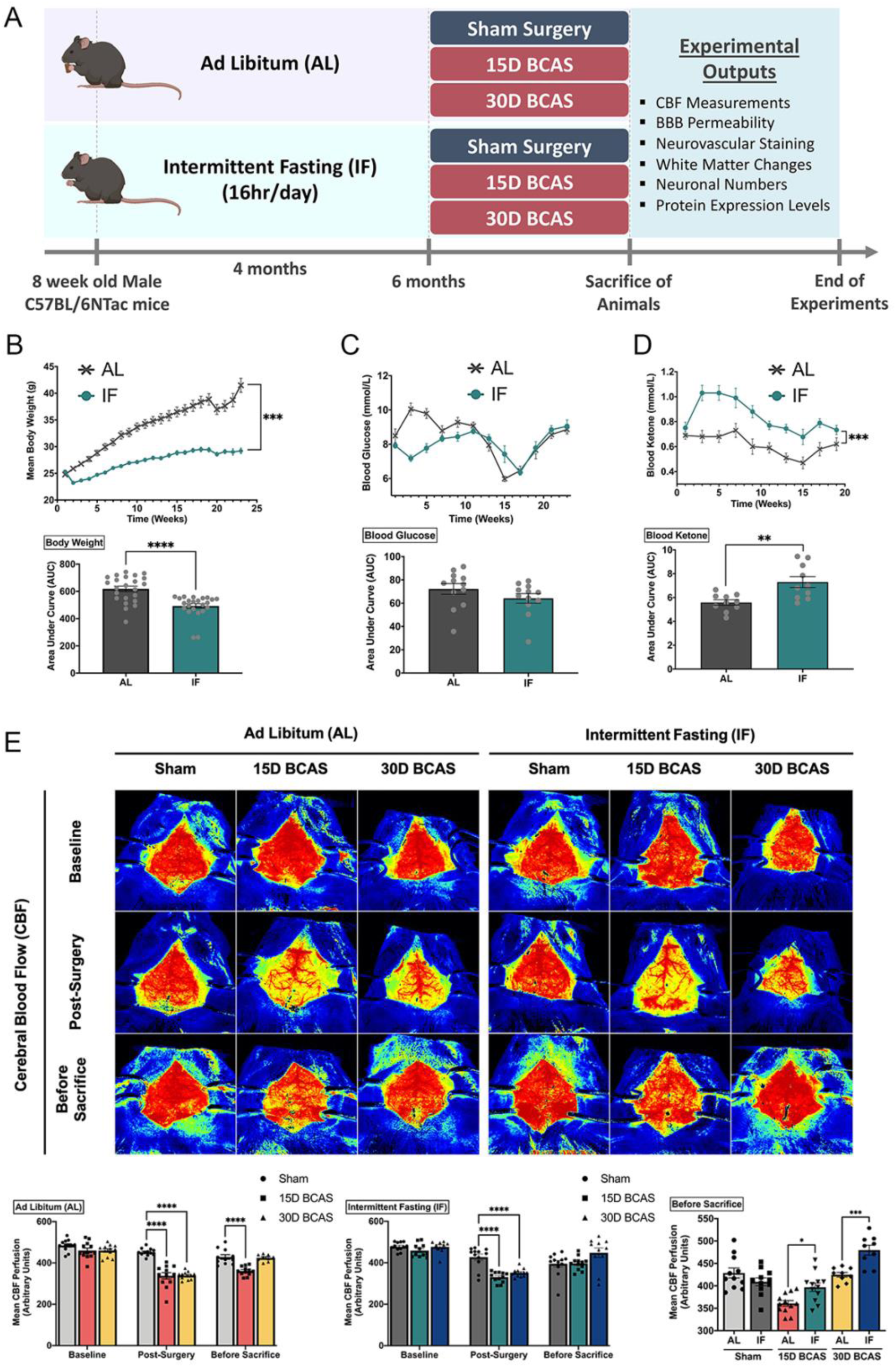
Effect of intermittent fasting on physiological measurements and blood flow in the brain following BCAS in a mouse model of VCI. **A**. Schematic representation of the experimental design and outputs performed. **B**. Mean body weight measurements over 25 weeks. ***P<0.001, ****P<0.0001 compared with AL. **C**. Blood glucose measurements over 25 weeks. **D**. Blood ketone measurements over 25 weeks. **P<0.01, ***P<0.001 compared with AL. **E**. Representative contrast images and quantification of cerebral blood flow at baseline, effective blood flow reduction post-surgery, and the final level of cerebral blood flow before sacrifice. The rate of blood flow was expressed in perfusion units (PU) using the PeriMed Software. *P<0.05, ***P<0.001 compared with corresponding AL BCAS; ****P<0.0001 compared with corresponding Sham. Data are represented as mean ± standard error of the mean of n=10-12 mice in each experimental group. Abbreviations: AL, ad libitum; BBB, blood brain barrier; BCAS, bilateral common carotid artery stenosis; CBF, cerebral blood flow; IF, intermittent fasting; VCI, vascular cognitive impairment.

### Bilateral Common Carotid Artery Stenosis (BCAS) Mouse Model

At six months of age, AL and IF experimental animals were further divided into three experimental groups: Sham, 15-day bilateral common carotid artery stenosis (BCAS) and 30-day BCAS. C57BL/6NTac mice were subjected to chronic cerebral hypoperfusion injury by bilateral common carotid artery stenosis (BCAS) surgery, as previously described (Shibata et al., 2004). Briefly, the animals were anesthetized with isoflurane (1.5-2% at a flow rate of 0.4-0.8 liter/min), and a vertical midline incision was made in the neck. The left and right common carotid arteries were isolated from surrounding tissues and exposed individually, and loosely ligated with silk thread for easy manipulation of the arteries. The common carotid arteries were each constricted with microcoils of an internal diameter 0.18mm, specifically designed for the mice (microcoil specifications: piano wire with gold plating, piano wire diameter 0.08mm, coiling pitch 0.5mm, and total length of 2.5mm; Sawane Spring Co. Ltd, Japan). The silk threads were removed, and the site of surgery was closed. The mice were monitored post-surgery until they were awake. Sham animals were subjected to a midline cervical incision and their common carotid arteries were exposed, but no microcoils were inserted. All animals were euthanised at their respective end points after BCAS for subsequent experimental analysis.

### Cerebral Blood Flow Measurements

The Laser Speckle Contrast Imager (PSI system, Perimed Inc.) was used to obtain high-resolution cerebral blood flow (CBF) measurements before BCAS (baseline), immediately after BCAS surgery (post-BCAS) and finally at the end points of BCAS (before sacrifice). As shown in **Figure 1E**, the brain regions of interest (ROI) between the bregma and the lambda were selected to measure arbitrary units of CBF in the area between the cerebral hemispheres. Briefly, the fur on the head was removed by shaving and the skull was exposed via a midline skin incision. The skull was cleaned gently using a cotton applicator with 1×phosphate-buffered solution (PBS). Throughout the imaging of CBF, the skull was kept moist and in order to improve imaging resolution, a non-toxic silicon oil was applied on the skull. Body temperature was maintained at 37 ± 0.5°C throughout the measurement periods. In the PeriCam PSI System, a CCD camera (2448×2048 pixels) that can take images in real-time was installed 10cm above the skull (speed 120 frames per second). Images were then acquired and analysed using a dedicated software package (PIMSoft, Perimed Inc.).

### Immunoblot Analysis

Animals were euthanized by inhaled carbon dioxide and the brains were harvested at their respective timepoints. Different brain regions (cortex, hippocampus, and cerebellum) were isolated immediately on ice and frozen in dry ice for future analysis (n=7 in each experimental group). Detailed immunoblot analysis procedures were performed as previously described (Fann et al., 2013). Briefly, brain tissues were homogenised in lysis buffer (Thermo Fisher Scientific, #78510) and combined with protease (Thermo Fisher Scientific, #78443) and phosphatase (Thermo Fisher Scientific, #78428) inhibitors to prevent proteolysis and dephosphorylation of proteins respectively during extraction, and then combined with 2x Laemelli buffer (Bio-Rad Laboratories, Inc., Hercules, CA, USA). Protein samples were then separated on 5-12.5% v/v sodium dodecyl sulfate (SDS) gel matrixes. Polyacrylamide gel electrophoresis of SDS-treated proteins were separated based on their sizes. The proteins on the gels were then transferred onto nitrocellulose membranes to allow for probing. The nitrocellulose membranes were incubated with the following primary antibodies: VEGF (Santa Cruz, sc7269), PDGFRβ (Cell Signaling, #3169), ZO-1 (Thermo Fisher, 61-7300), Occludin (Invitrogen, 711500), Claudin-5 (Thermo Fisher, 35-2500), JAM-A (Santa Cruz, sc25629), MBP (Cell Signaling, #78896S), Cleaved Caspase-3 (Cell Signaling, #9664), Total Caspase-3 (Cell Signaling, #9662), MMP2 (Cell Signaling, #87809S), MMP9 (Millipore, AB19016), MT1-MMP (Cell Signaling, #13130S), Malondialdehyde (Abcam, ab6463), Superoxide Dismutase (Abcam, ab13498), Glutathione (Abcam, ab19534), Vinculin (Cell Signaling, #13901) and β-actin (Sigma-Aldrich, A5441). Following primary antibody incubation overnight, membranes were washed for 10 minutes thrice with 1×TBST before incubation with horseradish peroxidase (HRP)-conjugated secondary antibodies (Goat Anti-Rabbit – Cell Signaling Technology, Danvers, MA, USA; Goat Anti-Mouse – Sigma-Aldrich, St. Louis, MO, USA) for 1 hour at room temperature with agitation. Following secondary antibody incubation, membranes were washed for 10 minutes thrice with 1×TBST. Finally, the membranes were imaged using ChemiDocXRS+ imaging system (Bio-Rad Laboratories, Inc., Hercules, CA, USA) after the substrate, enhanced chemiluminescence (ECL), was added (Bio-Rad Laboratories, Inc., Hercules, CA, USA). ImageJ software (Version 1.46; National Institute of Health, Bethesda, MD, USA) was used to quantify proteins in relation to their corresponding housekeeping gene (β-actin/Vinculin).

### DiI Vasculature Staining

At the end of each timepoint, a subset of animals (5 animals per experimental group) were deeply anaesthetized with isoflurane, and perfusion through the heart was performed as described previously (Li et al., 2008). Cardiac perfusion began with 25mL of chilled 1×PBS (pH 7.4) followed by perfusion of 10mL DiI solution (Sigma Aldrich, D-282, #42364, Invitrogen), and finally perfusion with 25mL of chilled fixative 4% paraformaldehyde. Once the perfusion was completed, the brains were harvested and placed in vials containing 4% paraformaldehyde solution for immersion-fixation overnight at 4°C. The brains were then embedded in 5% agarose before sectioning by a vibratome (Leica VT1200) at a 100μm thickness and collected on slides. The tissue sections were viewed under a confocal microscope under a Texas Red Fluorescence Filter (TissueGnostics, TissueFAXS Slide Scanner).

### Evans Blue Staining

At the end of each timepoint, a subset of animals (6-7 animals per experimental group) underwent Evans Blue (EB) Staining analysis as described previously (Wick et al., 2018). Briefly, the animals were anesthetised using isoflurane. 2% EB solution (Sigma Aldrich, E-2129-10G, diluted in 1xPBS and filtered, 2ml/kg) was injected through the femoral vein into the mice. The EB was allowed to circulate in the mice for 30mins. At the end of the 30mins, the mice were euthanized with carbon dioxide. Cardiac perfusion of 20ml 1×PBS was performed to remove residual EB in the blood, after which the brain was harvested and stored in dry ice. Brain tissues were weighed, and the amount of 50% trichloroacetic acid (TCA) solution (diluted in 0.9% saline) was calculated based on a 1:2 ratio of weight (mg):volume (μl). The tissue was homogenised in TCA solution. The mixture was then sonicated (10 cycles, 30 seconds on, 30 seconds off). Subsequently, the mixture was then allowed to incubate overnight at 4°C on a rotator to allow for the complete extraction of EB from the brain tissues. The TCA-lysate mixture was then centrifuged (30mins, 15,000rcf, 4°C) and the supernatant was collected. The supernatant was loaded in a 96-well plate with supplementation of 95% ethanol in a 1:3 ratio, respectively. The plate was then placed into a spectrophotometer at 620nm to determine the EB concentration.

### Luxol Fast Blue and Cresyl Violet Staining

Luxol fast blue (LFB) staining and expression of myelin basic protein (MBP) revealed myelin integrity, while cresyl violet staining and expression of cell death markers were indicative assessments for neuronal loss in the hippocampus and different brain regions. Mouse brain tissues were fixed in 10% neutral buffered formalin and then processed into paraffin wax blocks. Coronal sections (3μm thick) were obtained via microtome sectioning (Leica Biosystems, RM2255) and collected on slides (Trajan Scientific and Medical, Australia). Luxol Fast Blue (LFB) staining was performed to detect the severity of white matter lesions (WMLs) at 5 regions (corpus collosum paramedian, corpus callosum median, caudoputamen, internal capsule and optic tract) in the brain. Briefly, tissue sections were de-waxed and rehydrated before immersion into the LFB solution (Abcam, UK) at 37°C overnight. Excess staining was removed using 95% ethanol treatment followed by washing in deionized water. Gray and white matter differentiation was initiated with the treatment of 0.05% aqueous lithium carbonate (Abcam, UK) for 20s, followed by 70% ethanol until the nuclei was decolorized. Sections were then immersed in Cresyl Violet solution (Abcam, UK) for 5 mins and the excess staining washed in deionized water. The sections were then dehydrated in an ethanol gradient (70-100%), and finally cleared in xylene and mounted onto the slides (Fluoromount-G Mounting Medium, Invitrogen). Brightfield images were taken under ×60 magnification using an Olympus upright Fluorescence Microscope BX53. A WML severity index was calculated by giving each brain region a given grade of either: Grade 0 (normal), Grade 1 (disarrangement of nerve fibres), Grade 2 (formation of marked vacuoles), or Grade 3 (disappearance of myelinated fibres). The severity of the WMLs were scored by three blinded examiners and the number of neurons in CA1, CA2 and CA3 regions of the hippocampus were counted as previously described (Nishio et al., 2010).

### Statistical Analysis

Experimental data were analysed by GraphPad Prism 8.0 software (GraphPad Software, San Diego, CA, USA). All values were expressed as mean ± standard error of the mean. One-way analysis of variance was used, followed by a Bonferroni post-hoc test to determine significance between experimental groups. A P-value<0.05 was deemed to be statistically significant.

## Results

### Intermittent fasting reduces body weight and increases plasma ketone levels, while improving cerebral blood flow recovery following chronic cerebral hypoperfusion

The summarized study design includes the timing of interventions, and blood and tissue collection (**Fig. 1**). As shown in **Figure 1A**, at eight weeks of age, the mice were randomly assigned to either the *ad libitum* (AL) or the intermittent fasting (IF) diet regimen. Male C57BL/6N mice were fed a normal chow diet (on a caloric basis: 58% carbohydrate, 24% protein, and 18% fat). Mice were randomly assigned to Sham, 15-day BCAS and 30-day BCAS groups at six months of age, when surgeries were performed. First, we monitored the physiological effects of IF on these mice. The IF group had a significantly lower body weight, a non-significant reduction in blood glucose level and a significant increase in blood ketone level than AL mice (**Fig. 1B-D**). We also examined the recovery of cerebral blood flow (CBF) in both AL and IF animals over 15 and 30 days following BCAS (**Fig. 1E**). At baseline, animals from all experimental groups had high CBF. Following BCAS surgery (Post-surgery), AL and IF groups had a significant reduction in CBF compared to their Sham counterparts, which did not show any reduction in CBF. Before sacrifice, at 15days and 30 days post-surgery, CBF was restored in BCAS animals.

### Intermittent fasting decreases microvascular leakage and maintains blood brain barrier integrity following chronic cerebral hypoperfusion

We next analysed the effect of IF against vascular pathologies following CCH. Fluorescence imaging of vasculature staining indicated an increase in the average number of microvascular leaks at 30 days after BCAS in AL animals, while IF reduced the number of microvascular leaks (**Fig. 2A-B**). BBB permeability after CCH was found to be increased following 15 and 30 days of BCAS under AL conditions. In contrast, IF reduced BBB permeability in sham and BCAS mice (**Fig. 2C-D**). To further analyse the structural integrity of the BBB, we next examined the expression levels of TJ proteins between endothelial cells at the cortex **(Fig. 2E, F)**, hippocampus **(Fig. 2G**,**H)** and cerebellum (**Fig. 2I, J**). TJ protein zonula occludens (ZO-1) showed a non-significant reduction in expression levels following 15-day and 30-day of BCAS at the cortex, a significant reduction at both 15-day and 30-day timepoints at the hippocampus but little change at the cerebellum in AL animals. Conversely, IF increased expression of ZO-1 in the cortex at both 15-day and 30-day timepoints, and in the cerebellum at 15-days (**Fig. 2E-J**). Expression of occludin was non-significantly reduced following 15-day and 30-day BCAS at the cortex, hippocampus and cerebellum regions in AL animals, while IF increased occludin expression at only the 30-day timepoint in all three regions (**Fig. 2E-J**). Claudin-5 levels were maintained following 15-day BCAS, but increased following 30-day BCAS in the cortex and cerebellum. In the hippocampus, Claudin-5 levels showed a non-significant reduction in AL mice following BCAS. IF maintained Claudin-5 levels following 15-day and 30-day BCAS in all three regions. Junctional adhesion molecule (JAM-A) expression was reduced in AL animals following BCAS in the cortex but not the hippocampus or cerebellum. IF maintained JAM-A expression levels in all three regions following BCAS (**Fig. 2E-J**).

**Figure 2:**
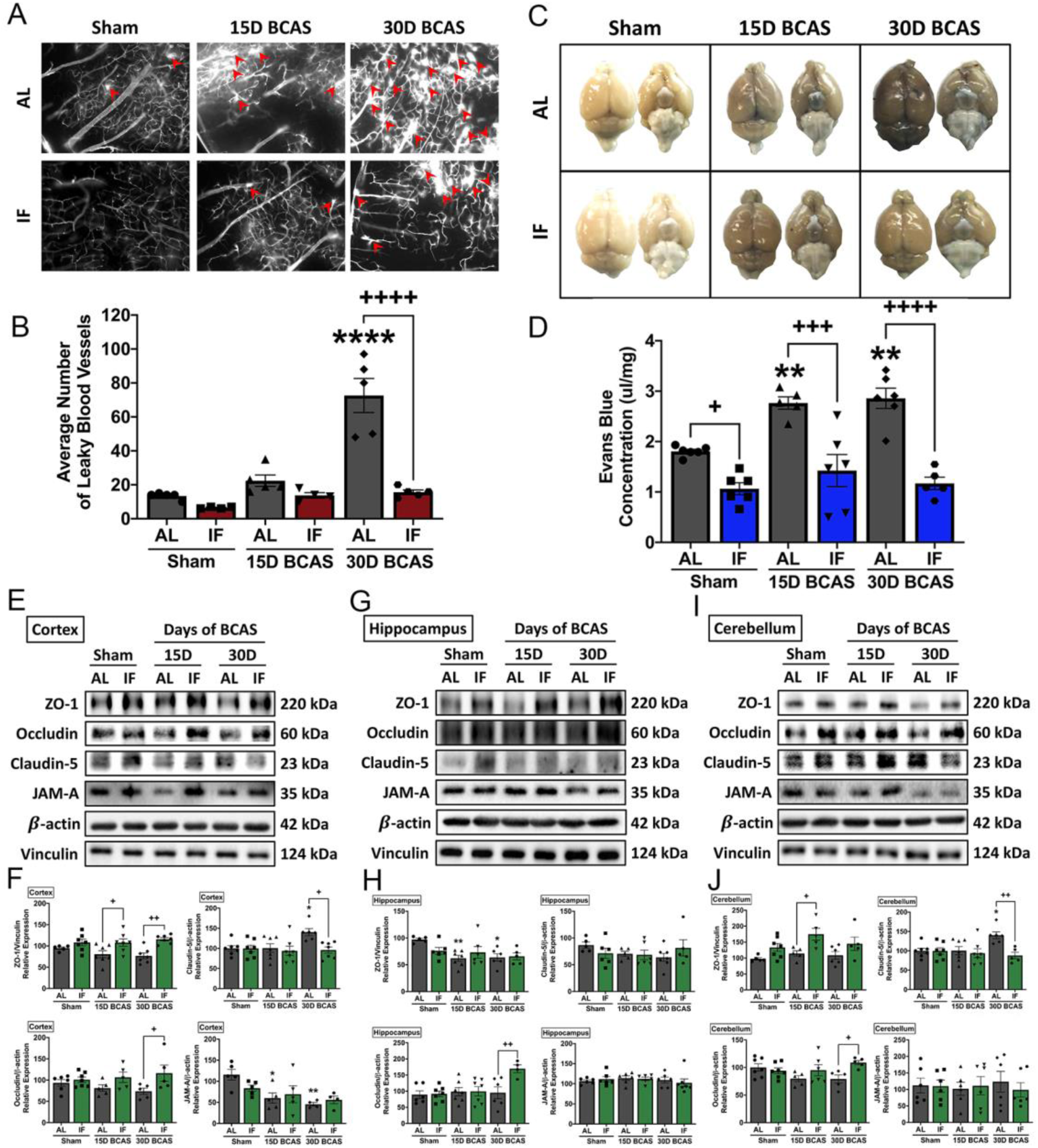
Effect of intermittent fasting on leaky blood vessel incidence and blood brain barrier integrity following BCAS. **A-B**. Representative DiI staining images and quantification illustrating leaky blood vessel incidence in AL and IF mice following BCAS. Data are represented as mean ± standard error of the mean of n=5 mice in each experimental group. ****P<0.0001 compared with AL Sham; ^++++^P<0.0001 compared with AL BCAS. **C-D**. Representative dorsal and ventral views of mouse brains injected with Evans Blue dye and respective quantification. Data are represented as mean ± standard error of the mean of n=5-7 mice in each experimental group. **P<0.01 compared with AL Sham; ^+^P<0.05, ^+++^P<0.001, ^++++^P<0.0001 compared to corresponding AL group. **E-J**. Representative immunoblots and quantification of tight junction proteins zonula occludens (ZO)-1, occludin, claudin-5 and junctional adhesion molecule (JAM)-A in the cortex **(E-F)**, hippocampus **(G-H)** and cerebellum **(I-J)**. Data are expressed as mean ± standard error of the mean. n=5-7 mice in each experimental group. β-actin or vinculin was used as a loading control. *P<0.05, **P<0.01 compared with AL Sham; ^+^P<0.05, ^++^P<0.01, compared with corresponding AL BCAS. Abbreviations: AL, ad libitum; BCAS, bilateral common carotid artery stenosis; IF, intermittent fasting; VCI, vascular cognitive impairment.

As cerebral vasculature pathology was evident following BCAS, we postulated involvement of vascular growth factors in BCAS-induced brain injury. Analysis of protein expression levels of vascular growth factors revealed a decrease in vascular endothelial growth factor (VEGF) in the cortex **(Fig. 3A, B)**, hippocampus **(Fig. 3C, D)** and cerebellum **(Fig. 3E**,**F)** following BCAS at 15-day and 30-day, while levels of platelet-derived growth factor receptor beta (PDGFRβ) were inconsistent. PDGFRβ was found to be decreased in the cortex, but not in the hippocampus or cerebellum under AL conditions (**Fig. 3A-F**). Interestingly, IF only decreased expression of PDGFRβ in the hippocampus following BCAS at 30-days.

**Figure 3:**
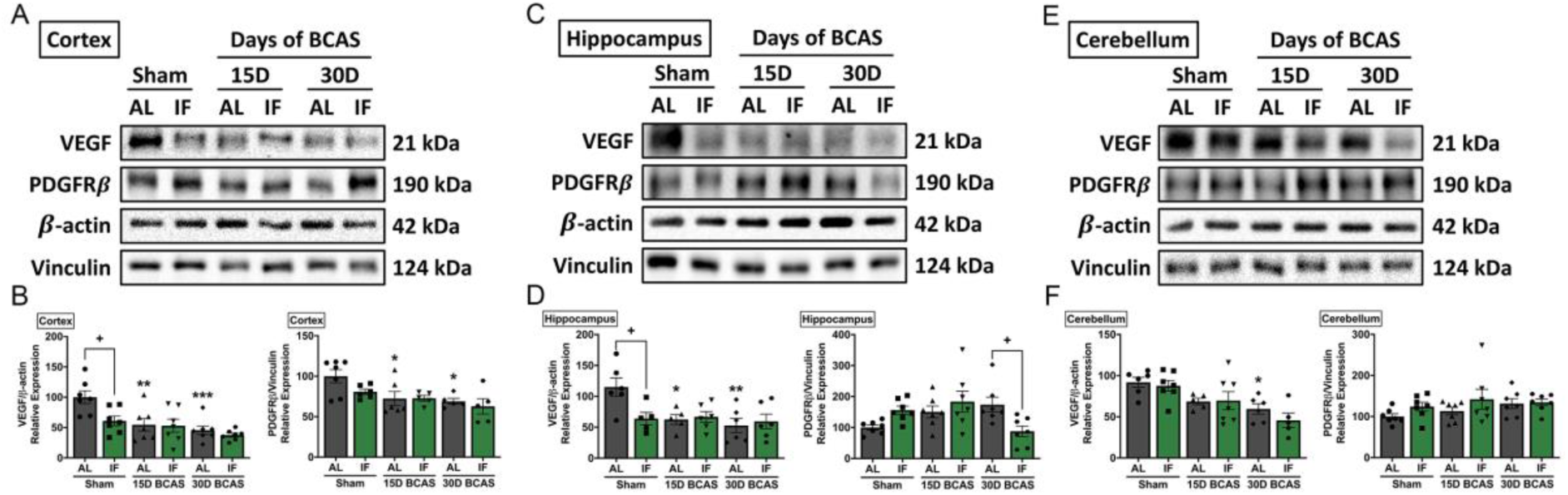
Effect of intermittent fasting on vascular endothelial growth factor and platelet derived growth factor receptor beta expression in the brain following BCAS. **A-F**. Representative immunoblots and quantification of vascular endothelial growth factor (VEGF) and platelet derived growth factor receptor beta (PDGFRβ) in the cortex **(A-B)**, hippocampus **(C-D)** and cerebellum **(E-F)**. Data are expressed as mean ± standard error of the mean. n=5-7 mice in each experimental group. β-actin or vinculin was used as a loading control. *P<0.05, **P<0.01, ***P<0.001 compared with AL Sham; ^+^P<0.05 compared with corresponding AL group. Abbreviations: AL, ad libitum; BCAS, bilateral common carotid artery stenosis; IF, intermittent fasting; VCI, vascular cognitive impairment.

### Intermittent fasting attenuates white matter damage and neuronal loss following chronic cerebral hypoperfusion

We next analysed the effect of IF on white matter integrity and neuronal loss. White matter integrity after 15 or 30 days of CCH was assessed in the corpus callosum (paramedian), corpus collosum (medial), caudoputamen, internal capsule and optic tract (**Fig. 4A-K**). White matter was disrupted in the corpus callosum (paramedian) (**Fig. 4B**,**C**) and caudoputamen (**Fig. 4F, G**) in AL mice following BCAS. At the corpus callosum (medial) (**Fig. D, E**) and internal capsule (**Fig. 4H, I**), AL animals showed a non-significant, but more severe disruption in white matter compared to their IF counterparts. We found no change in white matter disruption at the optic tract (**Fig. 4J, K**). We also found a decrease in expression of myelin basic protein (MBP) in the cortex but not hippocampus or cerebellum following 30-day of BCAS in AL animals (**Fig. 4L-Q**). IF increased MBP expression in the cortex and hippocampus in BCAS mice at 30-day (**Fig. 4L-M, N-O**).

**Figure 4:**
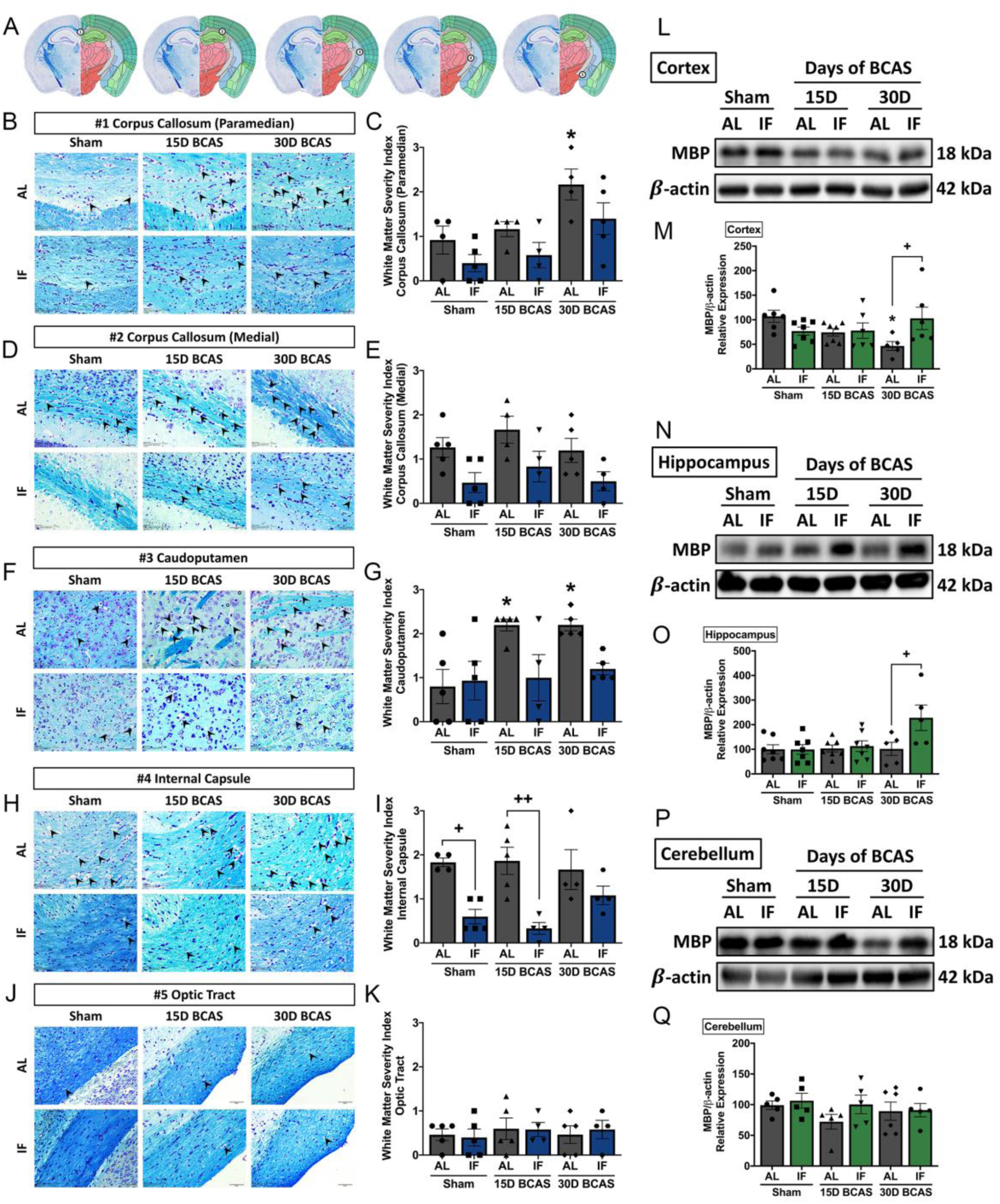
Effect of intermittent fasting on white matter integrity in the brain following BCAS. **A**. Schematic diagrams illustrating areas where white matter severity was measured, namely the corpus callosum (medial), corpus callosum (paramedian), caudoputamen, internal capsule and optic tract. **B-K**. Representative Luxol fast blue stained images and quantification illustrating white matter changes at the corpus callosum (paramedian) **(B-C)**, corpus callosum (medial) **(D-E)**, caudoputamen **(F-G)**, internal capsule **(H-I)** and optic tract **(J-K)**. The severity of white matter damage was graded as follows: Grade 0=no disruptions, Grade 1=disarrangement of nerve fibres, Grade 2=formation of marked vacuoles, and Grade 3=disappearance of myelinated fibres. Magnification x60. Scale bar, 20 μm. Images were taken under identical exposures and conditions. Data are represented as mean ± standard error of the mean. n=5 mice in each experimental group. *P<0.05 compared with AL Sham; ^+^P<0.05, ^++^P<0.01 compared with corresponding AL group. **L-Q**. Representative immunoblots and quantification of myelin basic protein (MBP) expression following BCAS in the cortex **(L-M)**, hippocampus **(N-O)** and cerebellum **(P-Q)**. Data are expressed as mean ± standard error of the mean. n=5-7 mice in each experimental group. β-actin was used as a loading control. *P<0.05 compared with AL Sham; ^+^P<0.05 compared with corresponding AL BCAS. Abbreviations: AL, ad libitum; BCAS, bilateral common carotid artery stenosis; IF, intermittent fasting; VCI, vascular cognitive impairment.

BCAS-induced neuronal loss was evident in hippocampal regions CA1 and CA3 (**Fig. 5A-F**). In the CA1 region, neuronal count was reduced at the 30-day timepoint in AL animals (**Fig.5A, B**). In the CA2 region, the neuronal count showed a non-significant reduction in neuronal numbers following BCAS in AL animals (**Fig. 5C, D**). In the CA3 region, there was a reduction in neuronal numbers at the 15-day timepoint following BCAS in AL animals (**Fig, 5E, F**). IF was able to rescue loss of neurons at the 15- and 30-day timepoints when compared to AL in all three regions (**Fig. 5A-F**). We next investigated if the loss of neurons were due to apoptotic programmed cell death following CCH (**Fig. 5G-L**). Our data showed that total caspase-3 levels remained unchanged in all three brain regions. Cleaved caspase-3 was found to show little change at the cortex (**Fig.5G, H**), a non-significant increase in the hippocampus (**Fig. 5I, J**), and a significant increase at the 15-day timepoint at the cerebellum (**Fig. 5K, L**) in AL animals, which suggests CCH-mediated neuronal loss to be partly mediated by activation of the apoptotic caspase pathway. IF animals had reduced expression of cleaved caspase-3 at the sham and 30-day BCAS timepoints at the cortex, and at the 15-day timepoint at the cerebellum.

**Figure 5:**
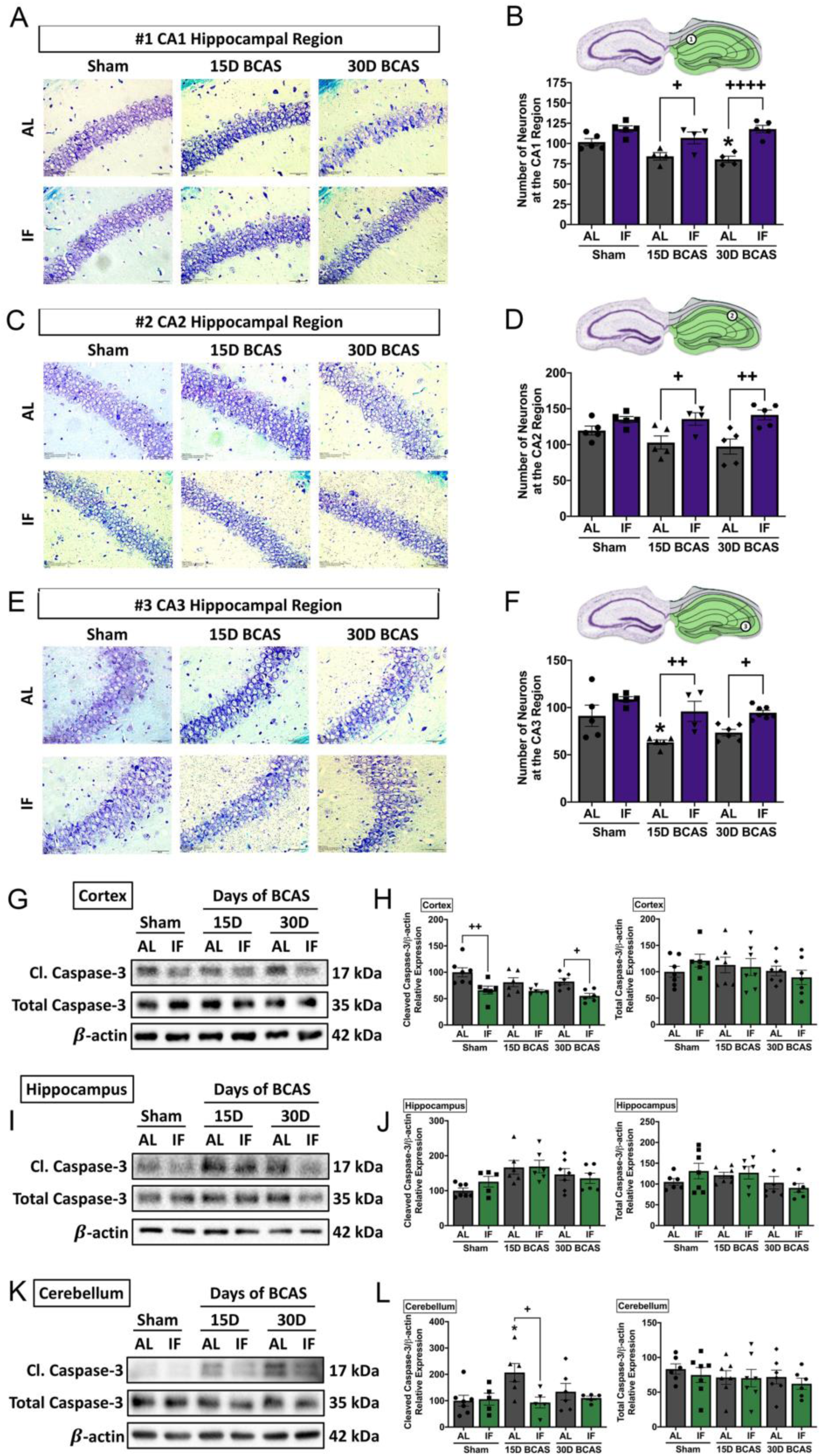
Effect of intermittent fasting on hippocampal neuronal loss and apoptotic death in the brain following BCAS. **A-F**. Representative cresyl violet images and quantification illustrating Nissl positively stained neurons in hippocampal CA1, CA2 and CA3 regions. Magnification x60. Scale bar, 20 μm. Images were taken under identical exposures and conditions. Data are represented as mean ± standard error of the mean. n=5 mice in each experimental group. *P<0.05 compared with AL Sham; ^+^P<0.05, ^++^P<0.01, ^++++^P<0.0001 compared with corresponding AL BCAS. **G-L**. Representative immunoblots and quantification illustrating apoptosis marker cleaved caspase-3 at the cortex **(G-H)**, hippocampus **(I-J)** and cerebellum **(K-L)**. Data are represented as mean ± standard error of the mean. n=5-7 mice in each experimental group. β-actin was used as a loading control. *P<0.05 compared with AL Sham; ^+^P<0.05, ^++^P<0.01 compared with corresponding AL group. Abbreviations: AL, ad libitum; BCAS, bilateral common carotid artery stenosis; Cl, cleaved; IF, intermittent fasting; VCI, vascular cognitive impairment.

### Intermittent fasting decreases matrix metalloproteinase and oxidative stress levels in the brain following chronic cerebral hypoperfusion

As various cellular and molecular pathways have been implicated in CCH-induced brain injury, we postulated the involvement of matrix metalloproteinases (MMPs) in the breakdown of the extracellular matrix in the brain that could mediate both vascular and neuronal pathologies. Our data established evidence for increased expression of membrane type-1 matrix metalloproteinase (MT1-MMP) at the cortex, and little change in the hippocampus and cerebellum following BCAS, while IF was able to maintain MT1-MMP levels at the 30-day timepoint in the hippocampus (**Fig. 6A-F**). Downstream pro-MMP-2 showed a non-significant increase following BCAS at the cortex, hippocampus and cerebellum in AL animals (**Fig. 6A-F**). IF was able to affect the CCH-induced increase of pro-MMP-2 levels at the cortex, hippocampus and cerebellum at the 30-day timepoint. However, we did not find any change in expression of pro-MMP-9 at the cortex, hippocampus and cerebellum, although pro-MMP-9 levels were downregulated in the hippocampus in IF BCAS animals at the 30-day timepoint (**Fig. 6A-F**).

**Figure 6:**
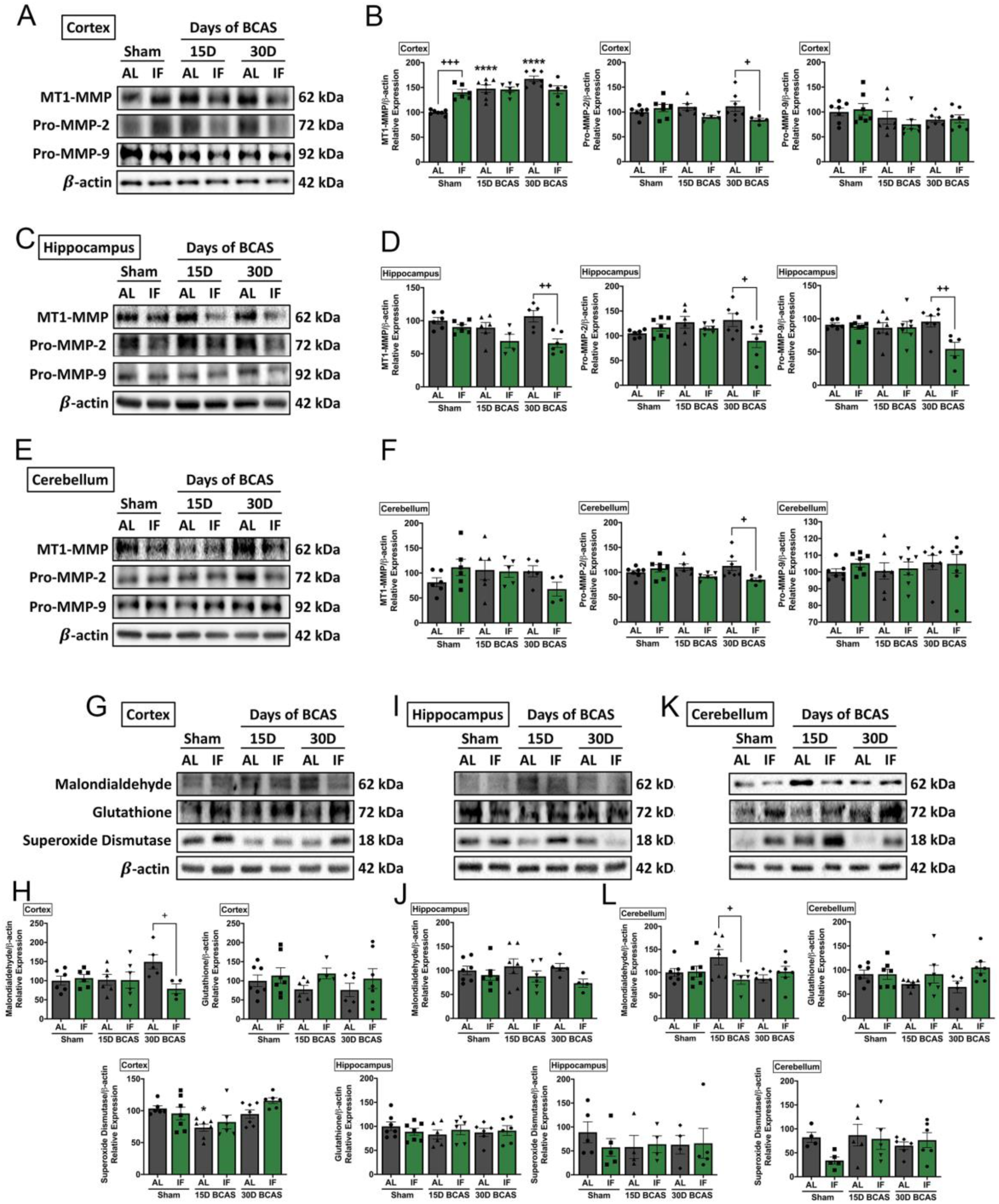
Effect of intermittent fasting on matrix metalloproteinase and oxidative stress in the brain following BCAS. **A-F**. Representative immunoblots and quantification illustrating matrix metalloproteinase (MMP)-2, its upstream membrane type (MT)-1 MMP (MT1MMP) and MMP-9 levels in the cortex **(A-B)**, hippocampus **(C-D)** and cerebellum **(E-F)**. Data are represented as mean ± standard error of the mean. n=5-7 mice in each experimental group. β-actin was used as a loading control. ****P<0.0001 compared with AL Sham; ^+^P<0.05, ^++^P<0.01, ^+++^P<0.001 compared with corresponding AL BCAS. **G-L**. Representative immunoblots and quantification illustrating oxidative stress marker, malondialdehyde, and anti-oxidant markers, glutathione and superoxide dismutase in the cortex **(G-H)**, hippocampus **(I-J)** and cerebellum **(K-L)**. Data are expressed as mean ± standard error of the mean. n=5-7 mice in each experimental group. β-actin was used as a loading control. *P<0.05 compared with AL Sham; ^+^P<0.05 compared with corresponding AL BCAS. Abbreviations: AL, ad libitum; BCAS, bilateral common carotid artery stenosis; IF, intermittent fasting; VCI, vascular cognitive impairment.

Another mechanism we postulated to be involved in CCH-induced brain injury was oxidative stress. We therefore analysed oxidative stress markers in the cortex (**Fig. 6G, H**), hippocampus (**Fig. 6I, J**) and cerebellum (**Fig. 6K, L**). Our data show that the oxidative stress marker, malondialdehyde, was non-significantly increased following BCAS injury in AL animals at the cortex, hippocampus and cerebellum. IF reduced malondialdehyde levels at the 30-day timepoint in the cortex and at the 15-day timepoint in the cerebellum (**Fig. 6G, H; 6K, L**). At the hippocampus, IF resulted in a non-significantly lower malondialdehyde level. Antioxidant glutathione levels were non-significantly lower in AL animals at the cortex and cerebellum following BCAS, and at the hippocampus, glutathione levels remained unchanged. IF was able to non-significantly maintain a higher level of glutathione as compared to its AL counterparts at the cortex and cerebellum, but showed no change at the hippocampus. Antioxidant superoxide dismutase was lower at the 15-day timepoint at the cortex but showed non-significant lower levels at the hippocampus and cerebellum (**Fig. 6G-L**).

## Discussion

Our findings indicate that CCH results in neurovascular pathology by activating MMP and promoting oxidative stress at the cellular level. Prophylactic IF attenuated microvascular leakage, BBB permeability, TJ breakdown, white matter injury, neuronal loss and cell death through a reduction in MMP activation and oxidative stress (**Fig. 7**). Collectively, these data are the first to demonstrate the effect of IF on vascular and neuronal pathologies following CCH injury, identifying IF as a potential therapeutic intervention for VCI.

**Figure 7:**
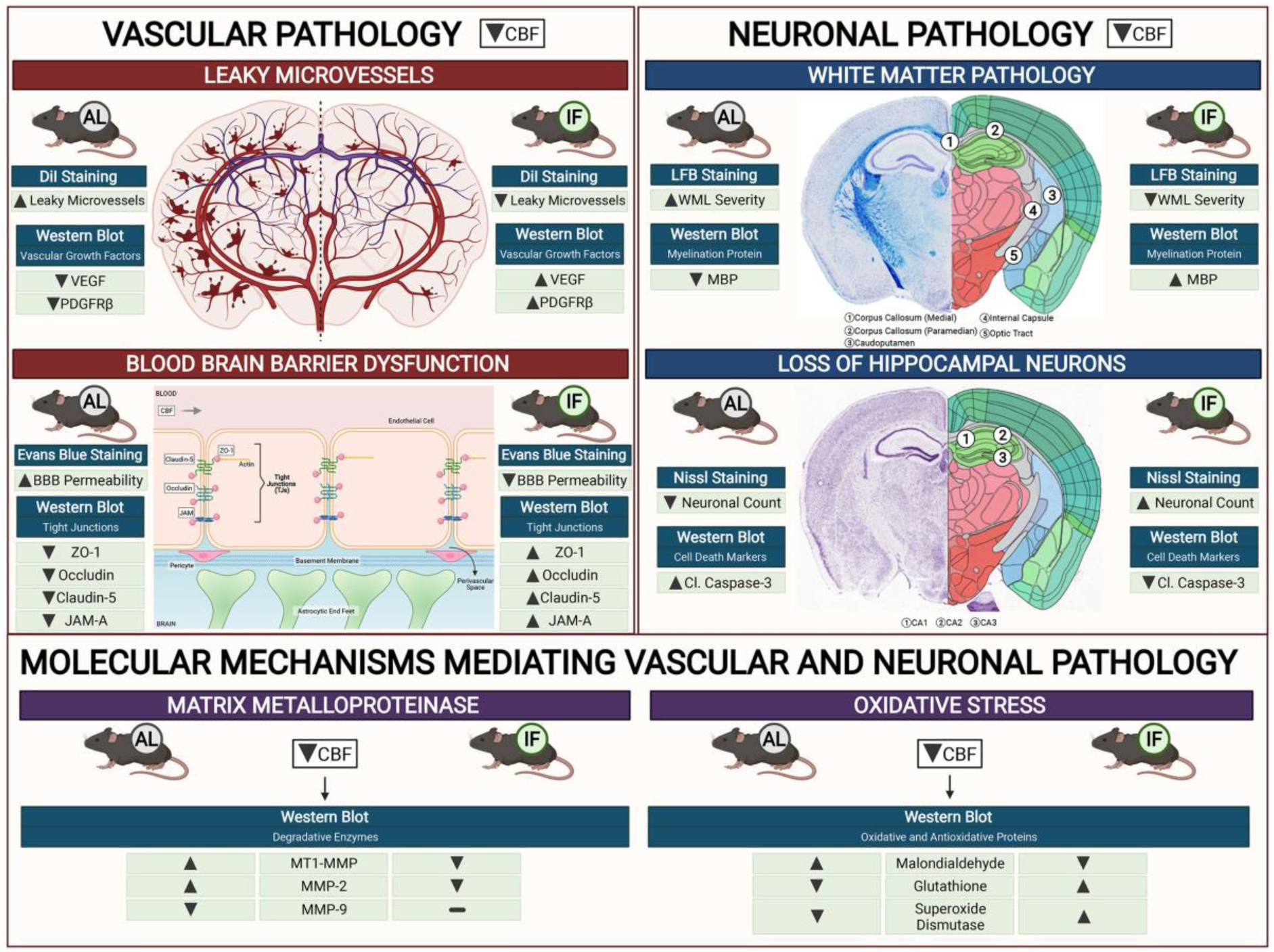
Schematic diagram illustrating the effects of intermittent fasting on the molecular mechanisms mediating vascular and neuronal pathology in the brain following BCAS in a mouse model of VCI. Vascular pathology in the brain following decreased cerebral blood flow (CBF) is defined as changes in the number of leaky microvessels and damage to the blood brain barrier (BBB). DiI staining was used to stain the vasculature, and IF was found to decrease the number of leaky microvessels in the brain following CCH. In addition, IF increased vascular growth factors in the brain, namely vascular endothelial growth factors (VEGF) and platelet derived growth factor receptor beta (PDGFRβ). Evans blue staining was used to assess the extent of BBB damage in the brain. IF was found to decrease BBB permeability through increased tight junction proteins zonula-occludens (ZO)-1, Occludin, Claudin-5 and junctional adhesion molecule (JAM)-A. Vascular pathology in the brain following decreased CBF is defined as changes in the white matter pathology and loss of hippocampal neurons. Luxol fast blue (LFB) staining method was used to stain and image 5 different white matter regions (corpus callosum medial, corpus callosum paramedian, caudoputamen, internal capsule and optic tract). IF decreased the severity of white matter damage in the brain following CCH, through maintenance of myelin basic protein (MBP) levels. Nissl staining was used to visualise the hippocampal neurons in the brain, and IF was found to be able to increase neuronal counts through reducing apoptotic death indicated by the levels of cleaved caspase-3 in the brain. IF was found to reduce membrane-type 1 MMP (MT1-MMP) and MMP2 levels in the brain following CCH. However, MMP9 was not involved in breaking down the extracellular matrix following CCH. IF reduced oxidative stress as evidenced through markers of lipid oxidation such as malondialdehyde, and antioxidative glutathione, and superoxide dismutase levels.

Following CCH, several hemodynamic changes occur in response to decreased blood flow in the brain. BCAS surgery induces a reduction of cerebral blood flow by approximately 30-40% before collateral development and recovery of blood flow (Nishio et al., 2010). Microscopic imaging of capillaries in the brain have revealed local heterogeneities in cortical blood flow supply during hypoperfusion, such as reduced red blood cell perfusion (Srinivasan et al., 2015). Our present data demonstrate that IF can alleviate vascular damage induced under hypoperfusion conditions, as IF animals had greater cerebral blood flow recovery at 15 and 30 days after BCAS surgery. Using a vasculature staining technique to observe vascular integrity, we found the number of microvascular leakages occurring due to CCH to be lower in IF than AL animals. Leaky microvasculature are structural abnormalities on small vessels that lead to reduced cerebral perfusion and are associated with aging and cognitive decline (Gregg et al., 2015; Martinez-Ramirez et al., 2014). Thus, it is plausible that IF maintained vascular integrity which improved CBF in the brain following hypoperfusion.

We investigated BBB permeability following CCH and the effects of IF on the integrity of the BBB in order to understand the mechanisms involved. TJ proteins such as ZO-1, Occludin, Claudin-5 and JAM-A act as molecular gates between endothelial cells of the BBB (Haseloff et. al., 2015). Increased BBB permeability has been reported in VCI patients and in animal models of CCH (Duncombe et. al., 2017). However, there were no previous studies of the effect of IF on BBB permeability. We found that IF was able to maintain the integrity of the BBB under hypoperfused conditions. Our data suggest that IF maintained BBB integrity via upregulating expression of TJ proteins. Our data are consisted with similar effects of IF on TJ proteins in the gut vasculature (Liu et. al., 2020). Following CCH, WMLs are reported to have a presumed vascular origin, and are associated with cognitive decline in the elderly (Murray et al., 2016). Reduced blood flow to the white matter regions have been shown to predict clinical development of WMLs in brain (Bernbaum et al., 2015). At the tissue level, breakdown of myelin sheaths causes myelin basic protein (MBP) network disassembly. A decrease in MBP expression with CCH therefore demonstrates structural disruption of the myelin sheath and hence formation of WMLs (Poh et al., 2020). We showed a striking difference between the severity of WMLs and MBP expression levels in animals under the AL and IF diets. We observed that IF animals subjected to BCAS exhibited less white matter injury and maintenance of MBP levels compared to those on AL diet. Similar results have been reported in multiple sclerosis mouse models where IF reduced demyelination in the spinal cord (Cignarella et. al., 2018).

Macroscopic pathological lesions due to CCH include the loss of neurons, particularly in the hippocampus which is associated with memory, behavioural changes and cognition (Cechetti et al., 2012). Sub-regions of the hippocampus – CA1, CA2 and CA3 are responsible for memory coding and retrieval. Specifically, the CA1 region is involved in the formation, consolidation, and retrieval of hippocampal-dependent memories (Bartsch et al., 2011); the CA2 region is involved in the formation of social memory (Chevaleyre & Piskorowski, 2016); and the CA3 region is involved in memory processes and neurodegeneration (Cherubini & Miles, 2015). Tightly orchestrated programmed cell death signalling events are activated in response to CCH, for which cell death markers such as caspase-3 play an important contributing role in apoptosis and hippocampal atrophy (Poh et al., 2020). We found that neuronal cell death, and cell death activation were present in AL animals following CCH, consistent with a previous study (Poh et al., 2020). In this study, we demonstrate less neuronal cell death in the CA1, CA2 and CA3 regions in IF mice following CCH. This implicates a mechanism through which IF improves cognitive functions. The cerebellum is also involved in cognition and behaviour of animals (Rapoport et. al., 2000). The increased caspase-3 levels observed in the cerebellum may also be closely linked to cognitive loss during CCH. IF attenuated cerebellar caspase-3 activation, which thus may mediate better cognitive performance.

It is important to elucidate the mechanism(s) at the molecular and cellular levels by which IF alleviates both vascular and neuronal pathologies following CCH. Here, we investigated pathways previously implicated in BBB breakdown, namely the activation of matrix metalloproteinases (MMPs) and oxidative stress. Previously, an increase in matrix metalloproteinases, particularly MMP-2 under CCH has been reported (Ihara et al., 2001; Kim et al., 2018). In our study, we showed a similar finding that MMP-2 was upregulated during CCH, and IF reduced the expression of MMP-2 in the brain. Similarly, there is also evidence that increased oxidative stress levels occur in the brain following CCH (Saxena et al., 2015). We observed that IF reduced CCH-induced increases in malondialdehyde levels, suggesting that reduced lipid peroxidation in the brain is a plausible mechanism through which IF attenuates neurovascular pathologies. Furthermore, IF increased anti-oxidant glutathione and superoxide dismutase levels in the brain following CCH. Our study suggests that IF exerts protective effects against vascular and neuronal pathologies through reduced levels of MMP and oxidative stress. In summary, we have provided substantial evidence that IF is effective in reducing vascular and neuronal pathologies following CCH. While our study establishes IF as a potential therapeutic approach for attenuating VCI-related pathologies, further research is needed to test its clinical potential.

## Declarations

### Ethics approval and consent to participate

All in vivo experimental procedures were approved by the National University of Singapore, Singapore Animal Care and Use Committee and performed according to the guidelines set forth by the National Advisory Committee for Laboratory Animal Research, Singapore.

### Availability of data and material

The datasets generated during and/or analysed during the current study are available from the corresponding author on reasonable request.

### Author contributions

Study conception and design: T.V.A., V.R., C.L.H.C., and D.Y.F.; experiment or data collection: V.R., and D.Y.F.,: data analysis: V.R., D.Y.F., and T.V.A.; data interpretation: V.R., T.V.A., and D.Y.F; writing-manuscript preparation and intellectual input: V.R., T.V.A., D.Y.F., Q.N.D., H.A.K., T.M.D.S., G.R.D., C.G.S., D-.G.J., M.K.P.L., and C.L-.H.C.,; supervision and administration: T.V.A., D.Y.F., M.K.P.L., and C.L-.H.C.

## Acknowledgements

All figures in this article were created using BioRender.

